# A balance between pro-inflammatory and pro-reparative macrophages is observed in regenerative D-MAPS

**DOI:** 10.1101/2022.08.17.504283

**Authors:** Yining Liu, Alejandra Suarez-Arnedo, Shamitha Shetty, Yaoying Wu, Michelle Schneider, Joel H. Collier, Tatiana Segura

## Abstract

Microporous annealed particle scaffolds (MAPS) are a new class of granular materials generated through the interlinking of tunable microgels, which produce an interconnected network of void space. These microgel building blocks can be designed with different mechanical or bio-active parameters to facilitate cell infiltration and modulate host response. Previously, changing the chirality of the microgel crosslinking peptides from L- to D-amino acids led to significant tissue regeneration and functional recovery in D-MAPS-treated cutaneous wounds. In this study, we investigated the immunomodulatory effect of D-MAPS in a subcutaneous implantation model. We uncovered how macrophages are the key antigen-presenting cells to uptake and present these biomaterials to the adaptive immune system. A robust linker-specific IgG2b/IgG1 response to D-MAPS was detected as early as 14 days post-implantation. The fine balance between pro-regenerative and pro-inflammatory macrophage phenotypes was observed in D-MAPS as an indicator for regenerative scaffolds. Our work offers valuable insights into the temporal cellular response to synthetic porous scaffolds and establishes a foundation for further optimization of immunomodulatory pro-regenerative outcomes.

## Introduction

Biomaterial scaffolds are broadly used in regenerative medicine to support the rebuilding of structural integrity, accelerate functional recovery, and serve as delivery depots. Traditionally, these scaffolds are designed to evade the immune system or suppress inflammation to mitigate rejection[1]. In recent years, the idea of immunomodulatory biomaterials has gained popularity. These material platforms can actively engage the complex immune system as part of their design. Leveraging the help of immune cells and the compounds these cells produce, immunomodulatory materials work collectively with innate and adaptive immunity in achieving an efficient and effective outcome ranging from better tissue repair to improved cell therapies or improved cancer therapies[2]. Given the complexity of eliciting the appropriate immune response at the right time, we first need to understand the local and systemic immune responses toward implanted materials.

During a typical host response to biomaterials, macrophages serve as both the first responders and the key mediators[1a]. Macrophage phenotype and involvement are both determinants in biocompatibility[3]. As antigen-presenting cells (APCs), macrophages bridge innate and adaptive immunity by surveying, internalizing, and presenting foreign signals to the adaptive immune cells during the inflammatory process[4]. A timely transition of macrophage phenotype from pro-inflammatory-biased to pro-regenerative dominant is needed to ensure the resolution of inflammation; however, a lingering pro-regenerative macrophage response leads to fibrosis and collagen deposition[5]. Therefore, if a biomaterial-driven, immune-mediated regenerative tissue response is desired, careful modulation of macrophage response is required [2c, 2d, 6]. Different macrophage phenotypes have been associated with divergent tissue responses, and conflicting observations arise due to many confounding factors (e.g., biomaterial design, animal models, characterization methods). For example, implanted extracellular matrix (ECM) scaffolds were shown to induce a pro-reparative macrophage response and led to constructive regeneration in a rat model[7]. On the other hand, a pro-angiogenic poly (2-hydroxyethyl methacrylate-co-methacrylic acid) hydrogel with 34 μm-pores had higher macrophage infiltration with a predominantly inflammatory marker expression (inducible nitric oxide synthase/iNOS and Interleukin 1 receptor type I) within a mouse implant model[8]. Adding to the complexity of macrophage phenotypes, another study with the same scaffold system demonstrated that this pro-angiogenic effect with minimal fibrotic response correlated with a higher number of infiltrating macrophages co-expressing both iNOS and CD206 (a pro-regenerative marker)[9]. Another ECM biomaterial system induced a pro-regenerative response for traumatic tissue injury in mice and a specific scaffold-associated macrophage (CD11b+F4/80+CD11c+/-CD206hiCD86+MHCII+) was identified[2a, 2b]. Our current understanding of macrophage-biomaterial interaction falls short of exploring macrophage phenotype change over time and using a combination of functional markers.

Microporous annealed particle scaffolds (MAPS) are a new class of granular hydrogels generated through the interlinking of spherical hydrogel microparticles (microgels), which produce an interconnected network of microgels that act as a porous scaffold[1d]. Previously, the treatment of skin wounds with MAPS in mice resulted in accelerated wound healing[1d] and regenerated skin[10] that was accompanied by a change in the CD11b immune response to the material. MAPS in this work were produced from spherical interlinked poly(ethylene glycol) microgel (70μm diameter, Supplementary figure 1), which were modified to contain the fibronectin minimal integrin binding sequence, RGD, and crosslinked with matrix metalloprotease (MMP) degradable peptides. Switching the amino acids at the site of MMP-mediated bond cleavage from L-to D-chirality induced rapid MAPS degradation and a pro-regenerative myeloid cell recruitment in wounds that was associated with to better skin regeneration (e.g., hair neogenesis and improved tensile strength)[2d]. Interestingly, *in vitro*, the change from L to D chirality resulted in slowed MMP degradation rates for D-chirality microgels (Supplementary figure 1) and D-peptide crosslinker was a poor activator of macrophage pathogen recognition receptors *in vitro*[2d]. Thus, the specific immune factors or mechanisms that enable D-MAPS to achieve a superior repair are still not fully uncovered.

Here, we show a comprehensive profile of the myeloid response to both L- and D-MAPS in a mouse subcutaneous implant model and explore the role of macrophages as key antigen-presenting cells (APCs) during biomaterial-tissue interactions. Advances in spectral flow cytometry and large multi-color panel designs enabled us to generate high-dimensional datasets and unexpectedly uncover the uptake of biomaterials in specific immune cells over time. We provide evidence that a balance between pro-regenerative and pro-inflammatory macrophage phenotypes is induced by D-peptide crosslinked scaffolds, which can polarize the scaffold microenvironment to a pro-regenerative innate and adaptive immune response. We demonstrate that a synthetic granular hydrogel system alone can recruit and mediate the accumulation of immune cells, specifically macrophages and other APCs.

## Results

### Inside D-MAPS: an early cytokine response correlated with better scaffold integration

To assess the immune response to MAPS, we injected four 50 μl L-or D-MAPS subcutaneously in C57BL/6 mice, respectively. After 4, 7, and 14 days, the implants were retrieved for histological analysis. Unlike a typical foreign body response where a collagen capsule is built up over time, there was minimal encapsulation around MAPS (Figure 1a). The general immune response towards both scaffolds was active but constructive, with an improved level of collagen deposition, granulation tissue and vascularization at 14 days post-implantation (Figure 1b,c,d). This is partly attributed to MAPS’s interconnected porous structure, which contains both openings on the surface for cells to enter and void space in-between the particles for cells to traverse (Supplementary figure 1). This porous nature of MAPS facilitates cellular infiltration into the scaffolds, regardless of the use of an L-or D-crosslinker (Figure 1a). The percentage of CD11b+ immune cells in D-MAPS was significantly higher than that in L-MAPS on day 4 and this trend was reversed on day 14. In both L- and D-MAPS, there was an accumulation of CD11c+ APCs on day 14 (Figure 1e,f). This indicated that MAPS actively recruited immune cells, and D-MAPS specifically induced an early CD11b+ immune cell accumulation. Strikingly, a drastically higher concentration of macrophage activation cytokines (IL-1β, TNF), key cytokine for Th2 immune response (IL-4), and anti-inflammatory cytokine (TGF-β) was also observed only in the D-MAPS on day 4 (Figure 1f)[11]. These results collectively suggested an early immune response in D-MAPS that called for a closer look.

**Figure 1:**
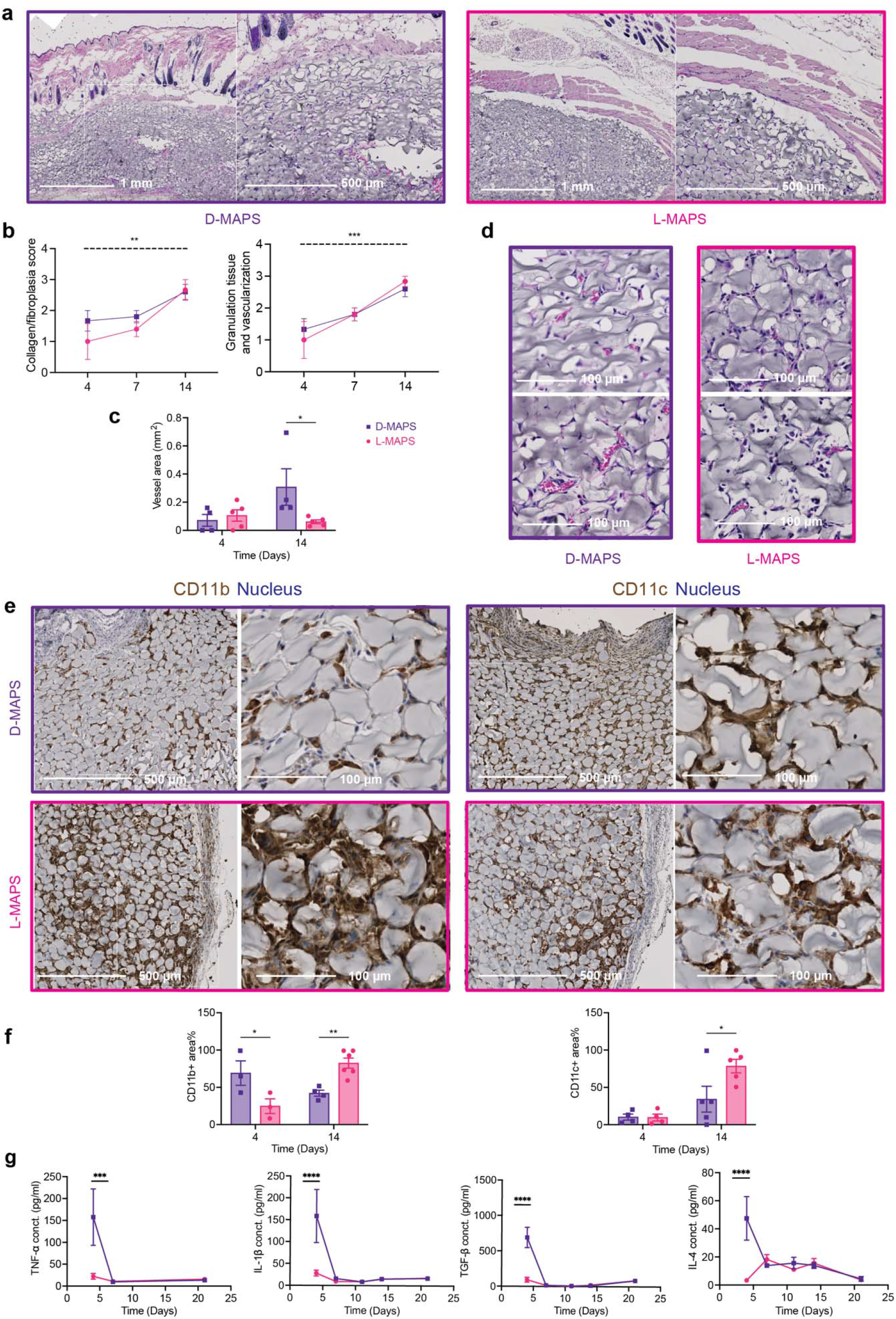
D-MAPS elicited an early immune response on day 4 and yielded a better implant integration outcome. **a**, representative pictures of Hematoxylin and eosin (H&E) staining on day 14. Each row from left to right showed pictures with objectives of 5x. **b**, histologic assessment of tissue integration level in L- and D-MAPS based on a modified 4-point scoring system established and agreed upon by two dermatopathologists. The dotted line indicated a significant time-dependent difference in both scaffolds. **c**, representative pictures of H&E staining at the center of implant with 20x objective. Scale bar, 100 μm. d, histologic assessment of angiogenesis in terms of blood vessel area per implant in L- and D-MAPS, measured in ImageJ. **e**, representative pictures of Immunohistochemical staining of CD11b and CD11c on day 14 at objectives 5x (skin/dorsal interface, scale bar, 500 μm), and 20x (capsule/ventral interface, scale bar, 100 μm). **f**, quantification of the percentage of CD11b+ and CD11c+ area among all the cell area. **g**, ELISA results of selected cytokine concentrations inside the hydrogel implants. Statistical analysis: two-way ANOVA with Šídák’s multiple comparisons test made between L- and D-MAPS groups only when there was a significance in the interaction term of scaffold type x time. ^*^ p < 0.05, ^**^ p < 0.001, ^***^p < 0.001,^****^ p < 0.0001. Error bars, mean ± s.e.m. n = 6 mice per group for a-f and n=5 for g with some data points removed due to experimental reasons. Experiments were repeated at least 2 times.

### Macrophages and FcεRI+ cells dominate immune cell infiltration into MAPS

With the help of multicolor flow cytometry, we set out to understand the complex immune infiltrate profiles inside L- and D-MAPS over the course of 21 days using an adapted panel from the literature[12]. This time window was chosen to explore the short-term immune response towards the materials: days 4 and 7 represent the early acute inflammatory phase. This is followed by a late inflammatory phase around 7 to 14 days, and the resolution phase occurs after 21 days (Figure 2a). The total infiltrated live cell number per implant weight and the percentage of CD45+ cells were similar between L- and D-scaffolds in the early stage (Figure 2b). L-MAPS retained a higher number of CD45+ immune cells on day 7 and day 14 while D-MAPS recruited more non-immune cells on days 14 and 21. This result corresponded with the improved tissue integration shown in the previous histology assessments. To get an overview of the cellular infiltration, a representative profile in both scaffolds was mapped out (Figure 2c). Macrophages and FcεRI+ cells dominated the immune cell infiltration and collectively they took up close to three-quarters of the total population across all time points. Macrophages and FcεRI+ (that can include Langerhan cells, monocytes, mast cells and other kinds of dendritic cells[13]) are the major immune modulators and APCs in the skin. Looking closer at the dynamics of each of the cell types, we observed that D-MAPS attracted a substantial number of eosinophils and dendritic cells at the early stage of host response (day 4) (Figure 2d), which might contribute to the early burst of cytokines in D-MAPS. A statistically significant increase in macrophage abundancy was observed in D-MAPS on day 14, which could be correlate with the initial cytokine response that included multiple macrophage activation factors (Figure 2d).

**Figure 2:**
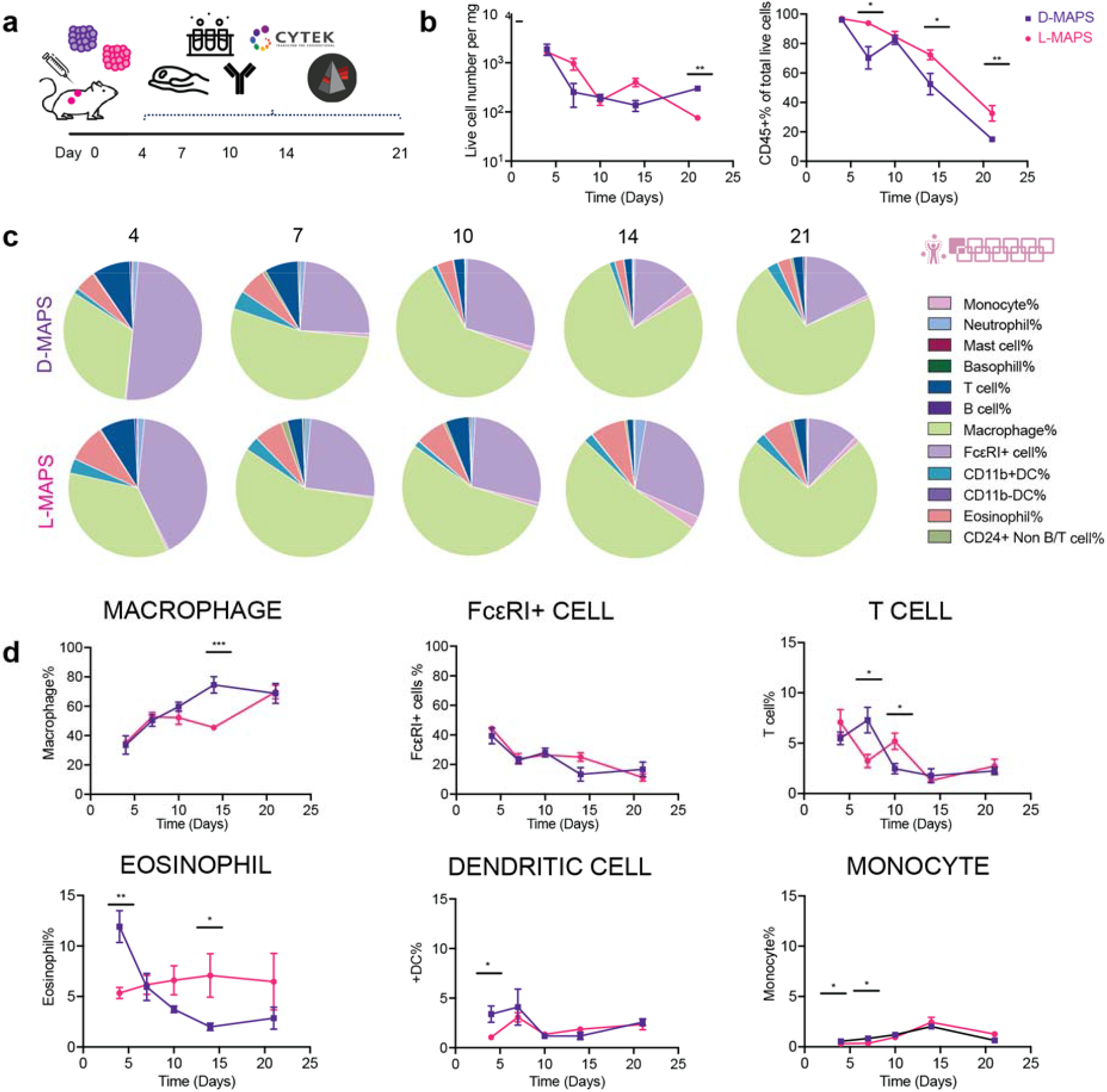
Macrophages and FcεRI+ cells dominated the immune cell infiltration. **a**, Scheme illustrating the experiment timeline. After the initial injections, implant extraction and flow cytometry were performed at designated time points (days 4, 7, 10, 14, 21). **b**, the total number of live cells (Zombie NIR-), the total number of live immune cells (CD45+) and their percentage among all live cells [monocyte (CD45+FceR1-MHCII-), neutrophil (CD45+Ly6G+), mast cells (CD45+CD117+FceR1-CD11b-CD11c-), basophil (CD45+CD117-FceR1-CD11b-CD11c-), T cell (CD45+MHCII-CD24-Ly6G-CD11b-CD11c-FceR1-) B cell (CD45+MHCII+CD24+Ly6G-CD11b-CD11c-FceR1-), macrophage (CD45+FceR1-MHCII±CD64+), FcεRI+ cells, CD11b+DC (CD45+Ly6G-FceR1-CD11b+MHCII+), CD11b-DC (CD45+Ly6G-FceR1-CD11b-MHCII+), Eosinophil (CD45+Ly6G-FceR1-CD11b+MHCII-) and CD24+ non B/T cell (CD45+CD24+MHCII-Ly6G-CD11b-CD11c-FceR1-)]. **c**, pie charts of myeloid cell abundancy across 5 time points for both D-MAPS and L-MAPS. Each number was an average of the results from 5 mice. **d**, macrophage, FcεRI+ cell, T cell, eosinophil, CD11b+ dendritic cell, monocyte percentages among all CD45+ live cells. Statistical analysis: two-way ANOVA with Šídák’s multiple comparisons test made between L- and D-MAPS groups only when there was a significance in the interaction term of scaffold type x time. ^*^ p < 0.05, ^**^ p < 0.001, ^***^p < 0.001. Error bars, mean ± s.e.m. n = 5 mice per group. The pink symbol in the middle right of the graph stands for the 11-color innate cell panel used in this figure.

### Pro-regenerative D-MAPS induced a balanced M1/M2 macrophage phenotype

To further dissect the infiltrated macrophage phenotypes inside MAPS, we designed a multicolor flow cytometry panel with six well-known macrophage functional markers and repeated the initial experiment[14]. CD86 and iNOS are the generally regarded “M1 pro-inflammatory markers” whereas CD206 and Arginase 1 (Arg1) are well-known “M2 pro-regenerative markers”[14]. MHCII and CD11c are established antigen-presenting cell markers, and when co-expressed with these two markers, CD86 also serves as the costimulatory signal necessary for T cell activation. Macrophages were defined as CD11b+F4/80+ live cells and their marker expression was traced over the initial 14 days post-implantation with L-or D-MAPS. This lineage definition covered most of the macrophage population inside the implants regardless of their origins (i.e., tissue-resident or monocyte-derived cells)[15]. Like our histology IHC results (Figure 1e, f), the CD11b+ cells were present at days 4, 7, and 14 through flow cytometry (Figure 3a). We found that these CD11b+ cells represented a significant fraction of the total live cell population, ∼60% at day 14 (Figure 3a). The initial influx of CD11b+ immune cells into D-MAPS was more pronounced than that in L-MAPS (Figure 3a). On day 4, almost all CD11b+ cells inside L-MAPS were F4/80+ macrophages, whereas D-MAPS also recruited other CD11b+ immune cells to facilitate the initial response (Figure 3b).

**Figure 3:**
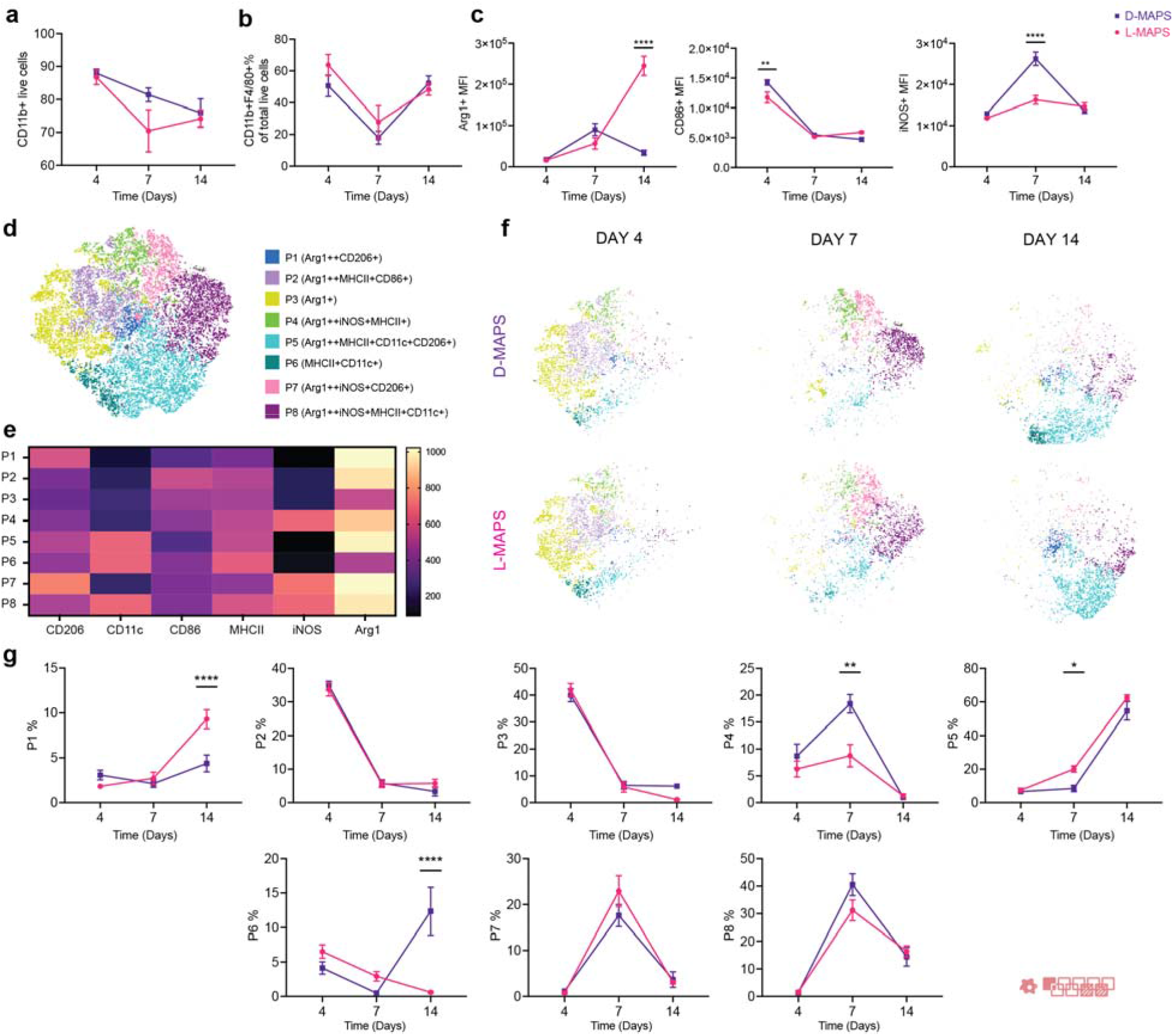
A balanced M1/M2 macrophage phenotype is induced in pro-regenerative D-MAPS. **a**, the percentage of CD11b+ cells in total live cells across 3 time points. **b**, F4/80+ macrophage percentage in CD11b+ live cells. **c**, MFI of Arg1, CD86, and iNOS in total macrophage population (CD11b+F4/80+) over time. **d**, tSNE clustering of the CD11b+ F4/80+ live cells and FlowSOM of 8 subpopulations at day 4, 7, 14 post-implantations. **e**. FlowSOM Heatmap of six phenotypical markers (Arg 1, CD206, CD11c, iNOS, CD86, MHCII) **f**. tSNE clustering of the CD11b+ F4/80+ live cells at day 4, 7, 14 post-implantations **g**. Percentage each sub-population from the total macrophage population at the specific time point. Statistical analysis: two-way ANOVA with Šídák’s multiple comparisons test made between L- and D-MAPS groups only when there was a significance in the interaction term of scaffold type x time. ^*^p < 0.05, ^**^p < 0.001, ^***^p < 0.001. Error bars, mean ± s.e.m. n = 6 mice per group. The red symbol in the bottom right corner of the graph stands for the 9-color macrophage panel used in this figure.

Overall, the phenotypes of the CD11b+F4/80+ population in both L- and D-MAPS were Arg1hiCD86hiMHCIIhi/+ at day 4 and Arg1hiCD206hi/+CD86+iNOS+/hiMHCIIhi at day 7, expressing both pro-repair and pro-inflammatory markers (Supplementary figure 3). At day 14, the general population shifted to a pro-regenerative, antigen-presenting profile (Arg1hiCD206+CD11c+MHCIIhi) (Supplementary figure 3). Macrophages in D-MAPS had a statistically higher CD86 level at day 4 and a significantly higher iNOS expression at day 7 (Figure 3c). By 14 days, the CD11b+F4/80+ population in L-MAPS expressed a drastically higher level of Arg1 than that in D-MAPS. At the designated inflammatory stage (days 4 and 7), D-MAPS promoted the pro-inflammatory marker expression, while at the resolution phase (day 14) it tamed down the pro-repair marker. Thus, the regenerative effect of D-MAPS was not associated with a particular pro-inflammatory or pro-regenerative macrophage phenotype but rather a balance of the two. Interestingly, macrophages in both L and D-MAPS through all days studied present a high expression of CD172a (Supplementary figure 3). CD172a is a glycoprotein found on the surface of myeloid cells, such as mast cells, granulocytes, dendritic cells (DCs), and macrophages, that have inhibitory effects in the immune response after the interaction with CD47[16].

To further characterize the macrophage phenotype profiles within L- and D-MAPS, we performed T-Stochastic Neighbor Embedding (tSNE) on the CD11b+ F4/80+ live cell population (Figure 3c). This algorithm creates two-dimensional visualization of high-dimensional flow cytometry data by calculating the distance between cell populations based on the expression of selected markers. We used the algorithm FlowSOM to guide the manual cell population gates on the spatially separated islands. These macrophage populations were characterized based on differential expression levels of each functional marker (shown in the legend as “++” indicating an MFI higher than 800, and “+’ indicating an MFI from 500 to 800 according to the FlowSOM heatmap). The tSNE/FlowSOM analysis showed that L- and D-MAPS attracted similar profiles of macrophages and polarized them into a broad range of phenotypes. A total of 8 distinct macrophage populations were identified. Population 1, which expressed have a higher expression of pro-regenerative markers Arg1 and CD206 was present in all 3 timepoints, increasing its proportion at day 14, mainly in L-MAPS. Notably, the proinflammatory markers CD86 and/or iNOS in some cases (population 2, 6, 7,8) were co-expressed with pro-regenerative markers CD206 and/or Arginase 1, indicating again that the CD11b+F4/80+ population is not binary pro-reparative or pro-inflammatory. Population 2 was highly expressed at day 4 (more than 30%) and expressed high levels of CD86 and MHCII, which showed a mature antigen-presenting function during the initial response to the implantation. At 7 days, Population 8 was the dominant population and had an M2-biased antigen-presenting phenotype (Arg1++ iNOS+MHCII+CD11c+). D-MAPS had a statistically significant increase in population 4 (Arg1++iNOS+MHCII+) at day 7 and Population 6 (MCHCII+CD11c+) at day 14. A shift toward these naïve macrophages may signify a change in the stage of inflammation and coincides with the increase in angiogenesis for D-MAPS at day-14. Further analysis will be needed to identify the phenotype of this CD11b+F4/80+ population. L-MAPS had a significant higher percent for Population 1 (Arg1++CD206+) at day 14. At day 14, most macrophage populations expressed pro-regenerative markers as the scaffolds became integrated and tolerated. D-MAPS retained a higher percentage of population 5 (Arg1++CD206+MHCII+CD11c+). Collectively, these observations confirmed that MAPS induced a complex macrophage response with unique expression profiles and a balanced M1/M2 phenotype was associated with the pro-regenerative D-MAPS.

### Infiltrated cells internalized MAPS throughout the implantation

When an injury or implantation occurs in the skin, antigen-presenting cells (e.g., macrophages, FcεRI+ cells) recognize and internalize danger signals to be presented to the adaptive immune system. In the case of MAPS implantation, we discovered using fluorescence microscope that the infiltrating cells degraded and ingested the hydrogel materials labeled with Alexa Fluor 647 (Figure 4a). The fluorescence of the internalized MAPS can also be detected as an intracellular marker using the new spectral flow cytometry technology. This allowed us to explore cell-driven material degradation and presentation across different time points. Both L- and D-MAPS implants contained a high number of MAPS+CD45+ immune cells across 21 days. The MAPS+% of total CD45+ cells peaked around day 7 to day 14 (Figure 4b). This showed an increased level of internalization by the infiltrated cells at the implants over time. Macrophages and FcεRI+ cells were the major populations infiltrating the scaffolds (Figure 2c) and also the main cells to internalize MAPS (Figure 4c). The higher level of L-MAPS internalization on days 4, 7, and 10 by macrophages and FcεRI+ cells correlated with the *in vitro* fast degradation profile of L-MAPS. On day 7, there was a significantly higher percentage of MAPS+ macrophages in D-MAPS than in L-MAPS (Figure 4c). A closer examination of macrophage phenotypes revealed that on day 4 population 7 (Arg 1++iNOS+C206+, population with both pro-regenerative and pro-inflammatory) internalized the most MAPS than population 3 (naïve, with a medium expression of Arg1) and population 6 (MHCII+CD11c+, antigen-presenting phenotype). At day 14, Both population 1 (Arg++CD206+) and population 5 (Arg++ MHCII+CD11c+ CD206+) internalized the most MAPS than population 3 and 6 (Figure 4e), regardless of the crosslinker used. This indicated the active M2-biassed antigen-presenting role macrophages played in the scaffold-induced immune response.

**Figure 4:**
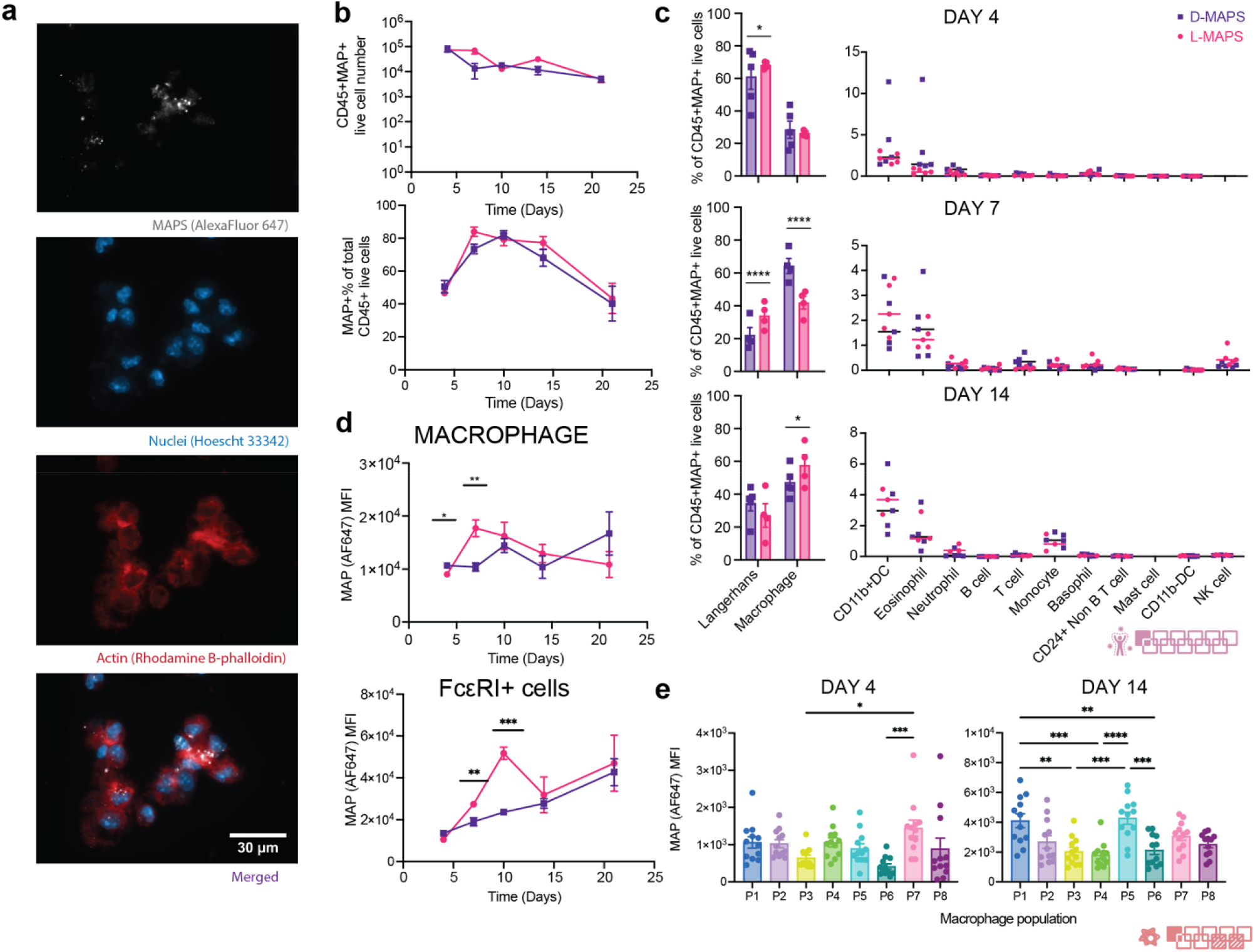
Infiltrated cells internalized MAPS throughout the implantation. **a**, Zeiss image of cells extracted from the implants showing MAPS internalization (scale bar, 30 μm); **b**, the total number of CD45+MAPS+ live cells and their percentage among all CD45+ live cells. **c**, MAPS+ immune cell abundance across 5 time points for both D- and L-MAPS. **d**, MFI of AF647 (ingested MAPS) in macrophage and FcεRI+ cell over time. **e**, MFI of AF647 (amount of internalized MAPS) in each macrophage combining results from both L- and D-MAPS at day 4 and 14. Day 7 was omitted in the plot since MFI AF647 expression was negligible for all populations. Statistical analysis: two-way ANOVA with Šídák’s multiple comparisons test made between L- and D-MAPS groups only when there was a significance in the interaction term of scaffold type x time. ^*^p < 0.05, ^**^p < 0.001, ^***^ p < 0.001. Error bars, mean ± s.e.m. n = 5 mice per group for b-d and n=15 for e, with some data points removed due to experimental reasons. The pink symbol in the middle right of the graph stands for the 11-color innate cell panel used in panel b-c of this figure. The red symbol in the bottom right corner of the graph stands for the 9-color macrophage panel used in this figure.

### MAPS+APCs migrated to draining lymph nodes and the spleen

APCs are essential players during the initiation and modulation of the adaptive immune response. These cells survey the body and uptake antigens. They then migrate to secondary lymphoid organs and present the peptide/MHC complexes directly to antigen-specific T cells. In the mouse, all DC subsets and some activated macrophages express the integrin CD11c, which is a key marker for APCs aside from MHCII. We hypothesized that some MAPS+ APCs migrated from the implants to draining lymph nodes (axillary and inguinal LN) and activated the adaptive immunity after seeing an increase of CD11c+% in dLN and the spleen (Supplementary Figure 4a). We adapted and designed a 20 color APC panel (OMIP-061) based on the literature to assess the heterogenous APC population[17]. On the organ level, we observed an increased level of MAPS+ signals in all the CD11c+ immune cells in the implants throughout the 14 days of implantation compared to baseline (Figure 5a). This trend shifted downwards as MAPS+CD11c+ cells migrated from the implant site toward secondary lymphatic organs (Figure 5a). There was an upward trend of MAPS+ signals in the draining lymph nodes (Figure 5a). On day 14, the migrating MAPS+CD45+ cells had passed from dLN to the spleen, resulting in a significantly higher level of MAPS+ above the baseline in the spleen (Figure 5a). Examining the phenotype of antigen-presenting cells in the implants revealed interesting transitions in cell types and phenotypes. The dominant cell type within the implants was macrophages (CD169+ macrophages and monocyte-derived macrophages/moM) on day 4 and day 7. This trend shifted over to dendritic cells from day 7 to day 14 (classical dendritic cells/cDC and monocyte-derived dendritic cells/mo-DC) (Figure 5b). cDC subsets recognize, process and present exogenous antigens to naive T cells and induce effective adaptive immunity[18]. More cDCs accumulated in L-MAPS on day 7 (Figure 5c). These cDCs in L-MAPS were derived from the residential DC population (Figure 5d) and biased towards a cDC2 phenotype (Figure 5e). cDC2 that are CD11b+F4/80+ followed same trends observed for macrophages in L-MAPS and D-MAPS. In D-MAPS, a higher percentage of cDCs remained undifferentiated (Figure 5e). pDCs are known as immunomodulating cells, regulating the immune response through type I interferon secretion, and participating in antiviral and pro-inflammatory responses[18]. A larger portion of CD45+ cells was categorized as pDCs in D-MAPS on day 7 (Figure 5c). These pDCs were expressing a higher level of CCR7 (trafficking), CD80 (co-stimulation), ICAM-1 (adhesion marker) (Figure 5f). An elevation of CCR7 expression in all the DC populations in D-MAPS on day 7 suggested a migratory phenotype after antigen uptake. These results mapped out APCs’ roles in bridging the local and systemic responses toward biomaterial implants by internalizing biomaterials and migrating to draining lymph nodes and the spleen.

**Figure 5:**
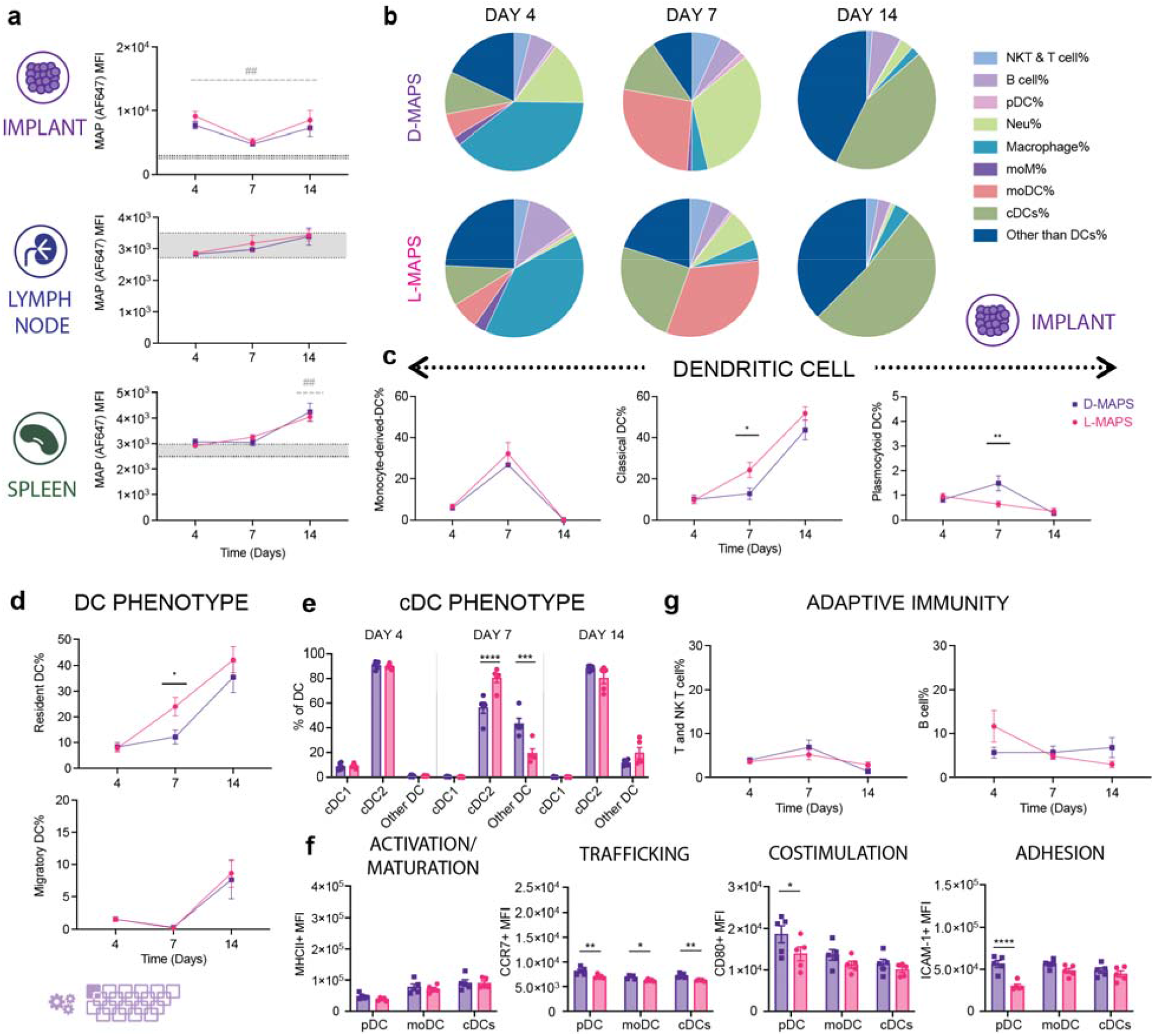
MAPS+CD11c+ cells migrated from the implant sites to draining lymph nodes and the spleen. **a**, MFI of AF647 among all CD11c+ live cells infiltrating the implants, residing in dLN or spleen. **b**, pie charts of APC cell abundancy across 3 time points for both D-MAPS and L-MAPS [NK and T cells (CD3+), B cells (pDCA-1-B220+), pDC (pDCA-1+ B220+), neutrophil (CD11b+ Ly6G+), macrophage (Ly6C+, CD169+, F4/80+), moDC, monocytes, and MCM (CD64+ CD11b+) monocyte-derived DC CD64-cDCs resident (CD11c^hi^ I-A/I-E^int^ CCR7^int^), cDCs migratory (CD11c^int^ I-A/I-E^hi^ CCR7^hi^), cDC1 (XCR1^hi^ CD172a^lo^) and cDC2 (XCR1lo CD172a^hi^)]. **c**, the percentages of dendritic cell subtypes cells among all CD45+ live cells infiltrating the implants across 3 different time points. **d**, the percentages of CCR7 high migratory DC and resident DC among all DCs on day 4, day 7 and day 14. **e**, the percentages of cDC1 and cDC2 among all DCs on day 4, day 7 and day 14. **f**, MFI of MHCII, CCR7, CD80 and ICAM-1 in pDC, moDC and cDC on day 7. **g**, the percentages of T and NK-T cells, B cells among all CD45+ live cells infiltrating the implants across 3 different time points. Statistical analysis: two-way ANOVA with Šídák’s multiple comparisons test made between L- and D-MAPS groups only when there was a significance in the interaction term of scaffold type x time. In panel a, after a two-way ANOVA, Dunnet method was used to compare the experiment groups with the baseline control group. ^*^ p<0.05, ^**^/## p<0.01, ^***^/### <0.001, ^****^/#### <0.0001. Asterisks stand for comparisons between L- and D-MAPS. Pound signs stand for comparisons between L-or D-MAPS and the baseline control (mice without implant). Error bars, mean ± s.e.m. n = 6 mice per group with some data points removed due to experimental reasons. The purple symbol in the bottom left corner of the graph stands for the 20-color APC panel used in this figure. pDC, plasmacytoid dendritic cell. moM, monocyte-derived macrophage. moDC, monocyte-derived dendritic cell (activation/maturation (MHCII), co-stimulation (CD80), and adhesion (ICAM-1)).

### A robust linker specific IgG2b/IgG1 response to D-MAPS was detected as early as 14 days post-implantation

Lastly, we examined whether the presentation of L-or D-MAPS by APCs to adaptive immunity in dLN and the spleen would lead to any adaptive immune response. An elevated total serum IgG concentration was detected on day 4 and day 7 for both L- and D-MAPS groups (Figure 6a), which was expected as an immediate systemic response to the scaffold implantation. Remarkably, by day 14, all mice with D-MAPS had developed linker-specific IgG response, whereas the response was less uniform in L-MAPS group, with some strongly responding mice and some non-responders (Figure 6a). A closer scrutinization of IgG subtypes revealed that D-MAPS induced an IgG2b-biased response with a similar level of IgG1 as L-MAPS group, while L-MAPS elicited a IgG1-dominant response. This observation is corroborated with the cytokine responses, with D-MAPS inducing broad range of cytokine secretion and L-MAPS primarily inducing IL-4 cytokine, a key cytokine for type 2 immunity. This also aligned with the APC phenotype profile in L-MAPS at day 7 since a cDC2-biased DC phenotype was closely tied to the Th2 pro-regenerative IgG1 response (Figure 5e). The B cell profiles in dLN and the spleen were drastically changed by material implantation (Figure 6b, Supplementary figure 3). An increase in the B cell population (B220+CD19+) on day 4 and day 7 was observed in dLN, which corresponded with the systemic serum IgG elevation (Supplementary figure 3). The percentage of CD95+ activated B cells went up in dLN on day 4 and day 7, and in the spleen on day 7. A similar trend was also observed for GL7+ germinal center B cells, with a peak on day 7 in both organs. Scaffold implantation also modulated T cell profile and increased both GATA3 and T-bet expression in dLN and the spleen (Supplementary figure 2). The evidence provided here further demonstrates that an adaptive immune response can be engaged and harnessed by synthetic pro-regenerative scaffolds[10]. We are actively investigating the implications of this observations, as recent report suggests materials-induced B cell responses are involved in wound healing.[19]

**Figure 6:**
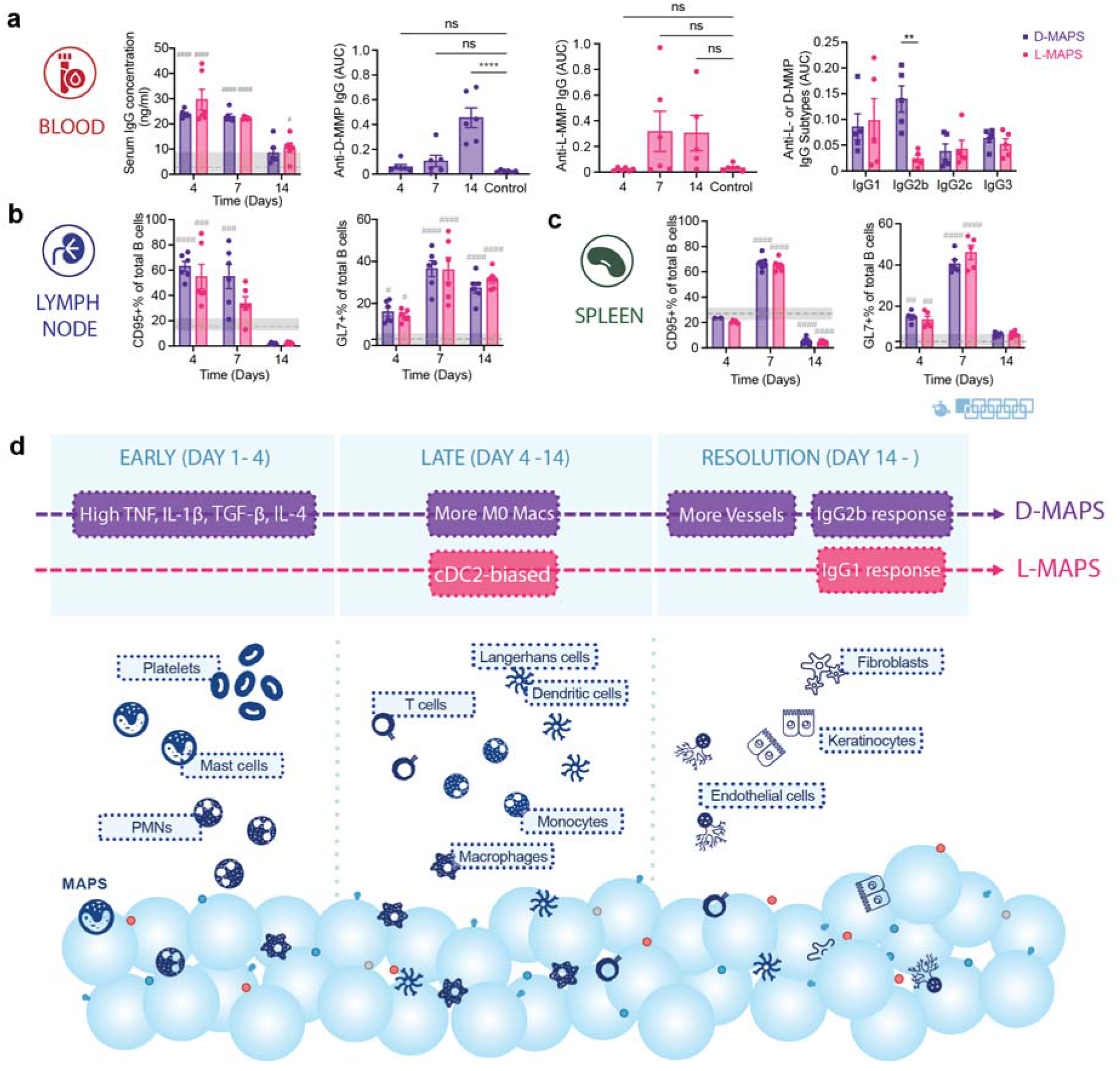
A robust linker-specific IgG response to D-MAPS was detected as early as 14 days post-implantation. **a**, ELISA results of total serum IgG concentration in the blood for mice with L-or D-MAPS implants, anti-D-MMP IgG level in mice with D-MAPS implants, anti-L-MMP IgG level in mice with L-MAPS implants and anti-L-or D-MMP IgG subtypes on day 14. **b**, CD95+% in total B cell population in draining lymph nodes (dLN) and spleen of mice with D or L-MAPS implants across 3 time points. **c**, GL7+% in total B cell population in draining lymph nodes (dLN) and spleen of mice with D or L-MAPS implants across 3 time points. **d**, Schematic illustration of the innate and adaptive immune response to D- and L-MAPS during subcutaneous implantation. The blue spheres represent MAPS, with peptides (colored circles and handles) on the surface. Statistical analysis: two-way ANOVA with Šídák’s multiple comparisons test made between L- and D-MAPS groups only when there was a significance in the interaction term of scaffold type x time. After a two-way ANOVA, Dunnet method was used to compare the experiment groups with the baseline control group. ^*^ p<0.05, ^**^/## p<0.01, ^***^/### <0.001, ^****^/#### <0.0001. Asterisks stand for comparisons between L- and D-MAPS. Pound signs stand for comparisons between L-or D-MAPS and the baseline control (mice without implant). Error bars, mean ± s.e.m. n = 6 mice per group. The blue symbol beneath panel c stands for the 9-color B panel used in panel b, c of this figure.

## Conclusions

Immunomodulatory biomaterials are versatile platforms to activate and direct specific elements of the immune system to treat conditions like cancer[2e, 2f, 20] and non-healing wounds[2d]. Different components of the biomaterial design, such as the peptide crosslinker in this study, can initiate and modulate the immune response. D-enantiomeric peptides carry important biological functions[21] and are widely used for their reduced proteolytic sensitivity[22]. Much evidence has shown that D peptides, when presented as part of a carrier platform like hydrogel scaffolds or nanoparticles, led to immune cell activation[23], controlled tissue inflammation[24], and peptide-specific adaptive immune response[10]. We previously showed that D-MAPS-treated skin wounds exhibited tissue regeneration accompanied with hair follicle formation, reduced scar formation, and improved mechanical stability[10]. D-MAPS generated D-enantiomeric peptide-specific innate and adaptive immune responses without the addition of cells, biological factors, or adjuvants[2d]. Wound regeneration with D-MAPS was linked to CD11b+ cells and an adaptive immune response. This manuscript first demonstrated that D-MAPS implant in the subcutaneous space had improved vascular scaffold integration, which we used as a measure of regenerative potential in a less complex environment than a wound. We confirmed the presence of CD11b+ cells and used this subcutaneous implantation model to analyze the immune cell profile via flow cytometry and ELISA. We found that D-MAPS had a statistically different cytokine profile early after implantation (day 4) (Figure 1g and Supplementary figure 2), including both pro-repair (IL-4 and TGF-β) as well as pro-inflammatory (IL-1β and TNF) cytokines. No difference in cytokines were observed at later timepoints, indicating that D-MAPS elicited a differential initial foreign body response. Overall, the immune foreign body response was similar between L and D-MAP as assessed with flow cytometry, showing higher expression of FcεRI+ cells and macrophages CD172a+. However, D-MAPS induced a linker specific antibody response in all animals tested, which was not the case for L-MAP (Figure 6) and L-MAP had a higher percentage of CD45% cells of the live cell population (Figure 2b). The macrophage phenotype for L-MAPS was significantly biased for Arg1+ at day 14, while D-MAPS was not, and had expression of both pro-repair and pro-inflammatory markers (Figure 3). The mode of biomaterial degradation is often thought to occur through cell released enzymes, however, we showed, for the first time, that MAPS were degraded through cellular internalization and cell-driven degradation, and that the internalized materials can be observed inside cells at the implant, lymph node, and spleen (Figure 5). These results showed that degradation occurs inside cells and that biomaterials are trafficked by cells to different cellular compartments. Additional studies using knockout mice for cells CD172a+ are required to determine whether the presence of those macrophages is associated, or the driver of the phenotype observed with both L and D-MAPS.

Activation of macrophages has been an active research area in tissue homeostasis, disease pathogenesis, and immune engineering[25]. Although an “M1 pro-inflammatory/M2 pro-reparative” dichotomy is often chosen for the simplicity of interpreting macrophage activation states and their functions[26], this naming method falls short of capturing the full picture[27]. Therefore, we used a combination of markers both in the assay and during the analysis to delineate and corroborate certain macrophage function[14]. Our findings suggested that a balanced macrophage phenotype is required at different stages of the immune response to pro-regenerative D-MAPS and their cooperation is needed to ensure an orderly tissue integration. Such findings are not possible without multidimensional and time-dependent analysis like the one demonstrated here.

## Author contributions

YL made substantial contributions to the design of the work, acquisition, analysis, and interpretation of data, as well as writing the manuscript. ASA performed the experiment and analyzed data that contributed substantially to the main figures. SS performed the experiment and analyzed data that contributed substantially to figure 6 and supplementary figure 2, 3. YW contribute in the design of B and T cell panel and data interpretation. MS oversaw and performed histology assessment that contributed substantially to figure 1. JHC contributed to data analysis and experimental design. All the authors discussed the results and contributed to writing portions of the manuscript and editing the manuscript. TS conceived the project, provided guidance and discussion throughout, and made substantial contributions to data analysis, figure preparation, and manuscript editing.

## Acknowledgements

We would like to thank the National Institutes of Health (R01AI152568) and all the members of the Segura lab and the Collier lab at Duke University for their support. Thank you to lab member Eleanor Caston for helping with part of the experiment. Thank you to Patrick Duncker, Ph.D. and Paul Dascani, Ph.D. at Cytek Biosciences for providing technical support on spectral flow cytometry and sharing insights on data interpretation. Thank you to Minerva Matos-Garner, MA, Ilana Palmer, MEd, Angus Bowers, MA, MSc, from Duke Engineering Graduate Communications and Intercultural Programs for their kind instruction and constructive suggestions on the manuscript. Thank you to Lindsay Riley, Ph.D. for her proofreading on the manuscript.

## Data availability

The data that support the findings of this study are available from the corresponding authors upon reasonable request. Source data are provided with this paper.

## Methods

### Microfluidic device design and fabrication

Microfluidic water-in-oil droplet generators were fabricated using soft lithography as previously described[1d]. Various devices were molded from the masters using Sylgard™ 184 PDMS (Dow Corning). The base and crosslinker were mixed at a 10:1 mass ratio, poured over the mold, and degassed under a vacuum for 20 minutes prior to curing overnight at 60 °C. Channels were sealed by treating the PDMS mold and a glass microscope slide with Hand-Held Tesla Coil at max volts for 15 seconds. After channel sealing, the device was left overnight at 60 °C to ensure the sealing efficiency. The channels were then functionalized by injecting 100 μl of a solution of Rain-X and reacting for 15 minutes at room temperature. The channels were then dried by vacuum and incubated overnight at 60 °C before use.

### Microgel generation and purification

Microfluidic devices and microgels were produced as previously described. Briefly, 20 kDa 8 arm PEG Vinylsulfone (JenKem Technology) was dissolved in 0.3 M triethanolamine (Sigma) pH 8.8 and pre-reacted with K-peptide (Ac-FKGGERCG-NH2, GenScript) and Q-peptide (Ac-NQEQVSPLGGERCG-NH2, GenScript) and RGD (Ac-RGDSPGERCG-NH2, GenScript) for at least one hour at 37°C. Then the pH was adjusted to around 4.5-5.5 to slow down the on-chip gelation and prevent any clogging at the flow-focusing region. The final precursor concentration should be 10% (w/v) 8 arm PEG Vinylsulfone with 1000 μM K-peptide, Q-peptide, and RGD. Concurrently, the cross-linker solution was prepared by dissolving the di-thiol matrix metalloproteinase sensitive peptide (Ac-GCRDGPQGIWGQDRCG-NH2, GenScript) in distilled water at 12 mM and reacted with 10 μM Alexa-Fluor 647-maleimide (Invitrogen) for 5 minutes. These solutions were filtered through a 0.22 μm sterile filter (Argos Technologies) before loading into 1 ml syringes (BD). The aqueous solutions did not mix until droplet segmentation on the microfluidic device. The pinching oil phase was a heavy mineral oil supplemented with 1% v/v Span-80 (VWR). Downstream of the segmentation region, a second oil inlet with a high concentration of Span-80 (5% v/v) and Triethylamine (3% v/v) was added and mixed to the flowing droplet emulsion.

These microgels were collected and allowed to gel overnight at room temperature to form microgels. The microgels were then purified by repeated washes with HEPES buffer (VWR), pH 8.3, containing 1% Antibiotic-Antimycotic (Invitrogen) and centrifugation. The purified microspheres were stored in HEPES buffer (pH 8.3 containing 1% Antibiotic-Antimycotic and 10 mM CaCl_2_) at 4 °C.

### Generation of scaffolds from microgels and Mechanical Testing

Fully swollen and equilibrated microgels were pelleted by centrifuging at 22 000 G for 5-20 minutes and discarding the supernatant till there is no more extra liquid to form a concentrated solution of microgels. 4 U/ml of thrombin (Sigma, 200 U/mL in 200 mM Tris-HCl, 150 mM NaCl, 20 mM CaCl2) and 10U/ml of Factor XIII (250 U/mL) were combined with the pelleted microgels and mixed via thorough pipetting before injection and allowed to incubate at 37°C for 30 minutes to form a solid hydrogel. Storage moduli of the hydrogel materials were determined by rotational rheometry (Anton-Parr, MCR301). The frequency sweeps were carried out at a shear frequency range from 10^-1 rad s-1 to 10^2 rad s-1 with a strain amplitude of 1%.

### Microgel Size and Void Volume Measurement

Microgel size was calculated from nine 4X Nikon Ti Eclipse pictures of three separate batches of microgels using a custom MATLAB code. For scaffold void volume measurement, three scaffolds were made for L- and D-MAPS (6 total). Using a Nikon Ti Eclipse mentioned above, 509 z-slices (0.275 μm each step) were taken in each gel at 40x, spanning a total distance of 140 μm. The images were analyzed for void volume fraction with IMARIS (Oxford instruments).

### Subcutaneous implantation

Subcutaneous implantation procedure was carried out in accordance with institutional and state guidelines and approved by the Duke University’s Division of Laboratory Animal Resources (DLAR) under protocol A085-21-04. Briefly, 7-12-week-old male and female C57BL/6 mice (n=5, all male for the first experiment and n=6, mixed gender for the repetition, Jackson Laboratory) were anesthetized with 3.0% veterinary isoflurane (Patterson) and maintained at 1.5-2.0% isoflurane. The hydrogels were loaded in a 1 cc syringe (EasyTouch) with a 29-gauge needle and each mouse received two or four 50 μl injections of the same type of scaffolds (L-or D-MAPS) on both flanks. After injection, mice were monitored until full recovery from the anesthesia. For baseline control, age-matched mice (n=7, mixed gender) with no implantation were used. All procedures were approved by the Duke University Institutional Animal Care and Use Committee and followed the NIH Guide for the Care and Use of Laboratory Animals.

### Sample extraction and flow cytometry

At designated time points, animals were euthanized and the blood, lymph nodes, the spleen and implants were collected. The implants from the same animal were pooled together, diced finely in 1 ml PBS and incubated on ice for 30 minutes to collect cytokine samples from the implant site. After spinning down and collecting the supernatant, the implants were enzymatically digested with the digestion solution (200 U/ml Collagenase IV and 125U/ml DNase I, both from Worthington Biochemical) in RPMI media (Invitrogen) for 15 minutes at 37°C. The resulting material was filtered through a 70 μm cell strainer (CELLTREAT) and washed once with 1x PBS to get a single cell suspension. These cells were then stained with Zombie NIR (BioLegend) for 15 minutes at room temperature to access the viability and blocked with Fcr Blocking Reagent (Miltenyi Biotec) for 10 minutes on ice, followed by surface marker staining for 30 minutes on ice. For intracellular marker staining, an intracellular fixation & permeabilization buffer set (Thermo Fisher) was used to prepare the samples. After staining, samples were washed and resuspended in 150 μl flow buffer (1x PBS, 1 mM EDTA, 0.2% BSA, 0.025% proclin) and analyzed on the Cytek NL-3000 Flow Cytometer. Data was acquired using SpectroFlo software and analyzed using FlowJo™ v10.8 Software (BD Life Sciences). Relative abundance of macrophage subpopulations was determined as a fraction of CD45+ viable cells (Gated first on scatter FSC x SSC, then doublet discrimination via FSC-A x FSC-H, prior to Viability Zombie NIR-/CD45+).

Clustering of flow cytometry data was completed by concatenating CD11b+ F4/80+ live cells pooled from all biological replicates and time points into one file and clustering with the tSNE (t-distributed stochastic neighbor embedding) plugin for 1000 iterations, operating at theta = 0.5, perplexityL=L50, and the Exact (vantage point tree) KNN algorithm, on six phenotypical markers (Arg 1, CD206, CD11c, iNOS, CD86, MHCII). Downstream clustering was performed with the FlowSOM algorithm, and the macrophages were phenotypically isolated by choosing 8 metaclusters. Each subset was further identified by the expression or absence of six phenotypical markers (Arg 1, CD206, CD11c, iNOS, CD86, MHCII).

### ELISA

To assess the anti-L-or anti-D-antibodies, sera collected from mice at designated time points (4-, 7- and 14-days post-implantation) were analyzed for antibody titers by ELISA. Briefly, plates were coated with L-or D-MMP peptide solution (20 μg /ml) or PBS overnight at 4 C. Plates were washed with PBS containing 0.05% Tween 20 (PBST) and then blocked with PBST containing 2% bovine serum albumin (PBST-BSA) for 1 hour at room temperature. Serum was serially diluted with PBST-BSA in 10-fold steps, applied to coated wells, and incubated for 2 hours at room temperature. To detect total IgG, HRP conjugated Fcγ fragment specific goat anti-mouse IgG (Jackson Immunoresearch) was used as the detection antibody and developed with TMB substrate (Thermo Fisher). For antibody isotyping, HRP-conjugated IgG subtype-specific, i.e., IgG1, IgG2b, IgG2c and IgG3, antibodies were utilized (Southern Biotech) in place of the total IgG detection antibody while all other steps were similar. A reported titer of 1 indicates no detectable signal above background. The optical density at 450 nm was read using a Spectramax i3X microplate reader (Softmax Pro 3.1 software; Molecular Devices).

All ELISA kits for quantifying different cytokine expression were purchased from Thermo Fisher Scientific and the experiment was performed according to the manufacturer’s protocol. The samples were tested without any further dilution. The optical density at 450 nm (measurement wavelength) and 570 nm (reference wavelength) were read using a Spark 20M multimode microplate reader (Sparkcontrol V2.3 software; Tecan).

### Histology analysis

At each time point, the implants were extracted, and one sample out of the four scaffold implants in the same mice was randomly selected for histology examination. For paraffin embedding, implant samples were fixed with 4% paraformaldehyde overnight at 4 °C before further processing. Paraffin blocks were sliced into 5□μm thickness for hematoxylin and eosin (H&E) staining and immunohistochemistry (IHC) staining.

For IHC staining, paraffin-embedded slides were deparaffinized with xylene and descendant ethanol, and then incubated in BLOXALL Blocking Solution (Vector Laboratories) for 10□minutes. After a wash in distilled water, the slides were incubated for 20Lminutes in 10□mM sodium citrate buffer with 0.05% Tween 20, pH□6 (VWR) at 95□°C using a microwave. The slides were brought to room temperature, rinsed in PBST (Phosphate Buffered Saline containing 0.05% Tween-20), and then incubated with Normal Horse Serum, 2.5% (Vector Laboratories) for 20 minutes at room temperature. These slides were then stained with rabbit primary antibody (anti-mouse CD11b Antibody from Novus Biologicals or anti-mouse CD11c Antibody from Cell signaling) at 4°C overnight. ImmPRESS® Horse Anti-Rabbit IgG PLUS Polymer Kit (Vector Laboratories) was then used for visualization of CD11b or CD11c in brown. Subsequently, the slides were washed in tap water, counterstained with Mayer hematoxylin solution (EMS), dehydrated in ethanol, and mounted with DPX (EMS).

For all the stained slides, full implant scans were performed using ZEISS Axio Scan.Z1. Images were quantified with an in-house algorithm in ImageJ to determine CD11b+ and CD11c+ cell area as a fraction of the total cell area. Briefly, the region of interest was drawn manually around the implant. The total cell area was calculated after color deconvolution with H&E DAB vector module and thresholding in the purple channel with ‘IJ_IsoData dark” method. Another thresholding method (“RenyiEntropy”) was implemented in the brown channel to quantify the positive area of CD11b or CD11c. The expression of each marker was calculated by dividing the positive marker area by the total cell area. For blood vessel area quantification, the total vessel area was calculated on the original H&E image by manually drawing regions of interest around the vessels.

H&E sections were also examined by a board-certified dermatopathologist (P.O.S.), who was blinded to the identity of the samples, to assess various aspects of implant integration. Granulation tissue formation and vascularization, collagen deposition and fibrosis/fibroplasia (early scar formation), and inflammation scores were evaluated by a modified 4-point scoring system[28] (listed in Tables S1–S3), which was established and agreed upon by two dermatopathologists.

### Statistics and reproducibility

All the statistical analysis was performed using Prism 9 (GraphPad, Inc.) software. Specifically, a one-way or a two-way ANOVA was used to determine the statistical significance. For one-way ANOVA, a post hoc analysis with the Dunnett’s multiple comparisons test was used. For two-way ANOVA, a multiple comparison analysis with the Šídák’s multiple comparisons test was used. When there was a baseline control in the groups, a multiple comparison analysis with the Dunnet method was used to compare the experiment groups with the baseline control group.

The subcutaneous implantation studies were repeated three times and the implants were examined with an 11-color innate flow cytometry panel or a 7-color macrophage panel. In the first experiment (n=5, all-male), we used a 7-color macrophage panel for the flow cytometry assay. The results indicated the same trend as the following studies and were included in the supplementary figures. In the second experiment (n=5, all-male), we used the 11-color innate panel for the flow cytometry assay. In the third experiment (n=6, mixed gender), we used the 9-color macrophage panel, 9-color B cell panel, 7-color T cell panel and 20-color APC panel for the flow cytometry assay. Samples with an excoriated skin layer or significantly lower live cell number due to technical issues were not included in the analysis. For the histological analysis, □L- and D-MAPS samples in B6 mice from two experiments were used (n□= □6 histological samples available out of an available n□= □10 implants). Implant samples that were injected superficially and with an excoriated skin layer were removed from the final dataset. For the cytokine and IgG assays, □L- and D-MAPS samples in B6 mice from two experiments were used (n□=□5, all-male for one study and n=6, mixed gender for another).

